# The trauma severity model: An ensemble machine learning approach to risk prediction

**DOI:** 10.1101/210575

**Authors:** Michael T. Gorczyca, Nicole C. Toscano, Julius D. Cheng

## Abstract

Statistical theory indicates that a flexible model can attain a lower generalization error than an inflexible model, provided that the setting is appropriate. This is highly relevant in the context of mortality risk prediction for trauma patients, as researchers have focused exclusively on the use of generalized linear models for risk prediction, and generalized linear models may be too inflexible to capture the potentially complex relationships in trauma data. Due to this, we propose a machine learning model, the Trauma Severity Model (TSM), for risk prediction. In order to validate TSM’s performance, this study compares TSM to three established risk prediction models: the Bayesian Logistic Injury Severity Score, the Harborview Assessment for Risk of Mortality, and the Trauma Mortality Prediction Model. Our results indicate that TSM has superior performance, and thereby provides improved risk prediction.

**Highlights:**

- We propose an ensemble machine learning model for trauma risk prediction.
- A hyper-parameter search scheme is proposed for model development.
- We compare our model to established models for trauma risk prediction.
- Our model improves over established models for each performance metric considered.

## 1. Background

Trauma is a global healthcare epidemic, accounting for 9.2% of all deaths and 10.9% of disability-adjusted life-years [1]. The potential impact of trauma injuries on one’s quality of life has inspired several studies on how we can improve the quality of trauma care, and consequently improve trauma patient outcomes. However, several of these studies require that we take a trauma patient’s injury severity (risk of mortality) into account, and there is no consensus as to which risk prediction model is most appropriate for use [2-9].

Interestingly, careful consideration of the methodologies used to develop these risk prediction models indicates that regardless of which model is most appropriate, there may be room for substantial improvement in the quality of risk prediction. One reason for this is that several risk prediction models have been developed from small data sets [2-5], which implies that these models may not represent the population appropriately [10]. Another reason is that every model developed from a large data set thus far has been a generalized linear model [6-9], and generalized linear models may be insufficient for capturing the potentially complex relationships that exist within trauma data.

For these reasons, our objective is to develop a risk prediction model from machine learning algorithms with data from the National Trauma Data Bank (NTDB) and to compare its performance to the performance of other established risk prediction models. This is achieved by comparing three established risk prediction models – the Bayesian Logistic Injury Severity Score (BLISS) [7], the Harborview Assessment for Risk of Mortality (HARM) [8], and the Trauma Mortality Prediction Model (TMPM) [9] – to a new machine learning model for risk prediction. This machine learning model is the Trauma Severity Model (TSM).

## 2. Methods

### 2.1. Data Summary and Processing

This study was performed using data from the NTDB for patients hospitalized in 2008, 2009, 2010, and 2012. The NTDB is currently the largest aggregation of trauma data in the United States and provides patient demographics, hospital demographics, ICD-9-CM diagnoses codes (ICD-9 codes), general trauma assessments, hospital identifiers, physiology values, and inhospital mortality [11]. The data set initially consisted of 2,865,867 patient records from 884 hospitals and 7,283 ICD-9 codes.

Risk prediction with this data set was formalized as a binary classification task. The input variables considered were patient demographics (age and gender), ICD-9 codes, and general trauma assessments (comorbidities, Glasgow Coma Scale response scores prescribed by a physician, injury mechanism, injury type, and intent of trauma). All input variables except age are treated as binary indicators specifying whether or not a patient had that particular condition. For Glasgow Coma Scale response scores, the eye response score is represented by 5 binary indicators (4 for response scores and 1 for not provided in the dataset), the verbal score is represented by 6 binary indicator variables (5 for response scores and 1 for not provided in the dataset), and the motor score is represented by 7 binary indicators (6 for response scores and 1 for not provided in the dataset). Response scores are treated as binary indicators to improve the performance of the linear models. The output variable is a binary indicator specifying whether or not the patient died prior to discharge from a hospital.

To ensure that model comparison is fair, we closely followed the data cleaning procedure from TMPM’s study, which is in accordance with the data cleaning procedures from BLISS and HARM’s studies. For patient selection, this involved excluding patients that had burns or an ICD-9 code unrelated to trauma (e.g., poisoning, drowning, or suffocation) (193,606), were admitted to a hospital that did not maintain complete documentation of relevant trauma diagnoses (655,440), were missing data (for age, comorbidities, gender, injury mechanism, injury type, intent of trauma, and outcome) (335,980), had pre-hospital mortality (60,234), were transferred to another hospital (848,885), were discharged to hospice care or another acute care hospital (16,429), withdrew care (18,395), or were less than one year old (47,693).

There are 2 differences between the patient selection process in this study and that in TMPM’s study. One difference is how we selected hospitals from which we selected patients. In TMPM’s study, the data set consisted of patients from hospitals that admitted at least 500 patients during at least 1 year of the study (hospitals with “substantial trauma experience”) [9]. We instead used all patients that were admitted to any hospital that kept complete records of all ICD-9 codes that were considered relevant in TMPM’s study. The reasoning for this is that some trauma centers that would qualify as having substantial trauma experience omitted relevant ICD-9 codes from their registry, and this could harm each model’s ability to provide accurate risk predictions [12]. Another difference is that TMPM’s study ensured complete documentation only for age, gender, and outcome when determining which patients to include. We extended this to also ensure complete documentation for comorbidities, injury mechanism, injury type, and intent of trauma. The reasoning for this is that (1) no additional patients were excluded because of these criteria, (2) this information is typically known at the time of admission and is relevant in determining patient outcome, and (3) no risk prediction model has ever given consideration to such a combination of variables.

Once the patient selection process was completed, an ICD-9 code combining procedure was performed, which followed the ICD-9 code combining procedure in TMPM’s study exactly (please see [9] for an overview of this procedure). This ICD-9 code cleaning procedure was followed by an additional pre-processing step that combined any ICD-9 code that appeared fewer than 5 times with the closest corresponding ICD-9 code (based on expert consensus). This consisted of combining a specific injury with a more general injury; an open injury with a closed injury; or a group of highly similar injuries that were poorly represented to one single injury. This additional pre-processing step improved the performance of all models in this study.

The patient selection process kept 1,385,795 patient records out of 2,865,867 patient records and 2,033 ICD-9 codes out of 7,283 ICD-9 codes. The ICD-9 code cleaning procedure from TMPM’s study collapsed these 2,033 ICD-9 codes into 1,272 binary indicators representing ICD-9 codes. Combining ICD-9 codes that appeared fewer than five times with what was determined to be the closest corresponding ICD-9 code collapsed these 1,272 binary indicators into 1,234 binary indicators. There are 74 other variables that represent patient demographics, general trauma assessments, and patient outcome, which leaves us with a sparse 1,385,795 by 1,308 matrix (all variables are binary indicators except age, which is numeric). Table 1 provides a brief summary of the demographics for this processed data set.

**Table 1:** Demographics for the processed data set.

| *Demographic Characteristics* |  |
| --- | --- |
| Age (Interquartile Range) | (23, 61) |
| Male (%) | 63.93 |
| Number of Hospitals* | 713 |
| Mortality before Discharge (%) | 3.77 |
| *Racial Characteristics* |  |
| White (%) | 64.52 |
| Black or African American (%) | 16.17 |
| Hispanic (%) | 12.74 |
| Asian (%) | 1.93 |
| Native American/Native Hawaiian/Pacific Islander (%) | 0.79 |
| Other (%) | 10.82 |
| Not Recorded (%) | 5.77 |

(\*) Specific hospital demographics not displayed due to this information changing each year in NTDB.
(||) Hispanic is denoted as an ethnicity in NTDB data, not race.

### 2.2. Experimental Setup

We considered two experiments for this study. The first experiment developed TSM, BLISS, HARM, and TMPM using information pertaining to ICD-9 codes in the processed data set as input variables (HARM and TMPM utilize dimensionality reduction procedures on ICD-9 codes). This first experiment will be referred to as the “ICD-9 experiment.” The second experiment developed TSM, BLISS, HARM, and TMPM using patient demographics (age and gender), information pertaining to ICD-9 codes, and general trauma assessments (comorbidities, Glasgow Coma Scale response scores, injury mechanism, injury type, and intent of trauma) in the processed data set as input variables. The second experiment will be referred to as the “augmented experiment.”

To ensure appropriate model development and assessment for these experiments, the processed data set was randomly divided into a training set for model development (60% of the entire data set), a validation set for optimizing model performance (20% of the entire data set), and a test set for model assessment (20% of the entire data set). Model development was performed using the h2o [13] and sandwich [14] packages in the R statistical software (Version 3.3.1) [15]. Optimizing model performance concerns minimizing log-loss (LL) on the validation set for these experiments [16].

### 2.3. Model Development

#### 2.3.1. BLISS

BLISS utilizes Bayesian logistic regression for risk prediction. To re-develop BLISS for this study, two different Bayesian logistic regression models for each experiment, where the prior distribution (a Laplace prior or a Gaussian prior) was varied. The model that had the lowest LL on the validation set was selected as BLISS for that experiment.

#### 2.3.2. HARM

HARM is a logistic regression model that takes advantage of the hierarchical structure of ICD-9 codes to reduce dimensionality. Specifically, ICD-9 codes and patient demographics are combined together to create new variables (based on expert consensus) that replace ICD-9 codes and patient demographics. These new variables are selected as the input variables for HARM using forward selection [16].

To develop HARM for this study, HARM’s variable combining procedure was followed as closely as possible with the processed data set – the NTDB does not account for diagnoses related to chronic obstructive pulmonary disease and ischemic heart disease, which correspond to three input variables in the original HARM model. Input variables in the data set that were not dealt with in HARM’s original study were still considered for the forward selection procedure, but these input variables were not combined with any other input variable. For the ICD-9 experiment, ICD-9 codes that were not replaced and new variables for ICD-9 codes were considered as input variables. For the augmented experiment, new variables for ICD-9 codes and patient demographics; ICD-9 codes and patient demographics not replaced; and general trauma assessments were considered as input variables. Forward selection was performed until LL on the validation set no longer improved for each experiment.

#### 2.3.3. TMPM

TMPM is a probit regression model that maps ICD-9 codes to numeric severity values (“MARC values”) in order to reduce dimensionality (please see [9] for an overview of TMPM’s model development). There is only one difference between our model development procedure for TMPM and that specified in its original study. TMPM was originally developed using information pertaining to a patient’s five largest MARC values as input variables. We instead used forward selection to select the input variables for TMPM. For the ICD-9 experiment, every MARC value a patient may have as well as first-order interactions between the five largest MARC values were considered for forward selection. For the augmented experiment, the same input variables in the ICD-9 experiment as well as patient demographics and general trauma assessments were considered for forward selection. Forward selection was performed until LL did not improve on the validation set. We found that this forward selection procedure improved LL of TMPM relative to following TMPM’s model development procedure exactly.

#### 2.3.4. TSM

TSM was developed using stacked generalization [17, 18]. Our approach to stacked generalization followed this sequence. First, several machine learning models, or base models, are created from four machine learning algorithms: logistic regression with the elastic net penalty [20], random forests [21], gradient boosted machines [22], and feed-forward neural networks [23]. The feed-forward neural networks were developed with the AdaDelta optimizer [24] and the Hogwild stochastic gradient update scheme [25]. During the training process, five-fold cross-validation was used to gather approximate out-of-sample risk predictions (cross-validated risk predictions) from each base model. Each base model’s cross-validated risk predictions are then combined to create a “meta-learner training set,” which is used to develop a higher-level model (a meta-learner). For clarity, the meta-learner training set consists of each base model’s cross-validated risk predictions as the input variables, and a binary indicator specifying whether or not the corresponding patient died prior to discharge as the output variable.

The meta-learner for TSM is a gradient boosted machine, which was developed using an exhaustive grid search where the only hyper-parameter varied was the maximum depth the trees in a gradient boosted machine were allowed to grow (from 1 to 16 with an increment of 1). All other hyper-parameters were set to their default values in h2o except the learning rate (which was set to 0.05), the annealing parameter for the learning rate (which was set to 0.99), and the number of trees developed (the default early stopping protocol in h2o was used with the validation set to determine how many trees to develop) [25]. The gradient boosted machine with the lowest LL on the validation set was selected as the meta-learner for TSM.

#### 2.3.5. TSM Hyper-Parameter Search Procedure

A benefit of using stacked generalization with cross-validation is that the meta-learner for TSM is developed from the same patients used to develop its base models. This allows appropriate comparison between TSM’s base models, TSM’s meta-learner, BLISS, HARM, and TMPM. However, in order to ensure appropriate model comparison, strong performing base models must be developed, which depends on the configuration of the hyper-parameter space for a hyper-parameter search procedure. This can be problematic in practice, as a hyper-parameter space is user-defined, and the user may configure the hyper-parameter space inappropriately [26]. To avoid this potential issue, we propose the following search procedure for hyper-parameter optimization.

First, a manual search is performed to determine an initial hyper-parameter space configuration for a machine learning algorithm. Then, machine learning models are developed using a random search for hyper-parameters within this initial configuration [27]. After 5 models are developed from this initial hyper-parameter space (10 for neural networks due to its larger number of hyper-parameters), a checking procedure is performed. For clarity when describing the sequence of this checking procedure, hyper-parameters denote the inputs of a machine learning algorithm, input values denote the hyper-parameters used to develop a machine learning model, and hyper-parameter interval denotes a dimension of a hyper-parameter space. (1) The top 2 performing models are selected based on their LL on the validation set (3 for neural networks). (2) The input values of these selected models are examined to determine where they lie on their corresponding hyper-parameter intervals. (3) For every hyper-parameter interval where the corresponding input values of all selected models are in the top (or bottom) quarter of their hyper-parameter intervals, shift these hyper-parameter intervals such that the top (or bottom) quarter of these original hyper-parameter intervals now represents the bottom (or top) quarter of new hyper-parameter intervals. If none of the hyper-parameter intervals shift after 5 (10) models are developed, then this checking procedure is performed after each subsequent model is developed and the checking procedure will give consideration to all models developed in this hyper-parameter space. If a hyper-parameter interval does shift, then the checking procedure is not performed until 5 (10) new models are developed, and the checking procedure will only give consideration to the models developed in this new hyper-parameter space. For this study, this search procedure was performed until 40 models were developed from each machine learning algorithm except neural networks, from which 80 models were developed. Table 2 provides the initial hyper-parameter space configuration of each algorithm.

**Table 2:**
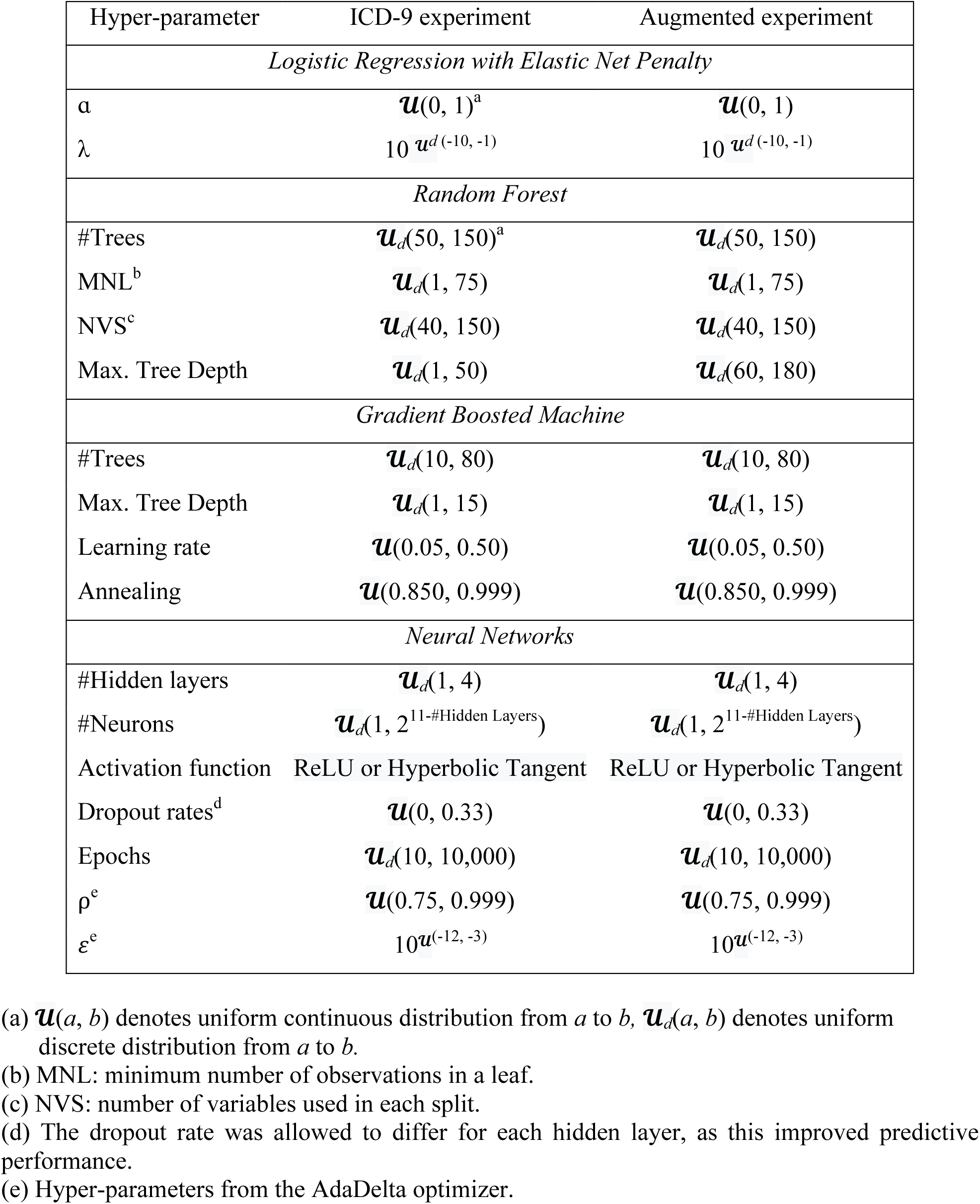
Initial hyper-parameter space configured for each machine learning algorithm. The number of trees in a random forest was not allowed to shift during our hyper-parameter search procedure.

#### 2.3.6. Machine Learning Model Calibration

Naively assessing the probabilistic calibration of each model in this study may be problematic, as non-linear machine learning models (random forests, gradient boosted machines, and neural networks) can have poor probabilistic calibration when the outcome event is rare, which Table 1 indicates [28, 29]. To avoid this potential issue, a balanced training set was developed for creating these non-linear models (all non-linear base models and meta-learner for TSM). This involved randomly over-sampling patients in the training set until the balanced training set was approximately 5 times the size of the original training set, and the number of patients who survived care was approximately the same as the number of patients who did not survive care [30]. Further, the base model from each non-linear algorithm that had the lowest LL on the validation set as well as the meta-learner for TSM were re-calibrated using isotonic regression before assessing their performance on the test set [28]. These isotonic regression models were developed with the prediction outputs of these models on the validation set.

### 2.4. Model Assessment

The performance of the established risk prediction models; TSM; and the logistic regression model developed with the elastic net penalty, random forest, gradient boosted machine, and neural network in TSM’s ensemble that has the lowest log-loss on the validation set (selected base models) are evaluated with six performance metrics, which may be divided into three groups: threshold metrics, rank metrics, and probabilistic calibration metrics (calibration metrics). The threshold metrics are classification accuracy (ACC) and F-score (FSC). These metrics are computed based on whether or not a risk prediction is above a user-specified threshold value. ACC and FSC range from 0 to 1, where larger values indicate better performance. A threshold of 0.5 was used when computing these metrics [31].

The rank metrics used in this study are the area under the receiver operating characteristic curve (ROC) [32] and the area under the precision-recall curve (APR) [33]. Rank metrics depend on the ordering of outcomes, and not the actual risk predictions. Provided that this ordering is preserved, the range of a model’s risk predictions does not affect its rank metric. These metrics measure how well positive cases (survival) are ordered before negative cases (mortality) and can be viewed as a summary of model performance across all possible thresholds. The ROC statistic may range from approximately 0.5 to 1, and APR may range from 0 to 1. Larger values indicate better performance.

Calibration metrics assess how well a risk prediction corresponds to a patient’s true risk of mortality. The calibration metrics considered in this study are log-loss (LL) [16] and the Hosmer-Lemeshow statistic (HL) [34]. For these metrics, smaller values indicate better performance, where 0 represents perfect probabilistic calibration.

In addition to assessing the models with these performance metrics, calibration curves [34] and precision-recall curves [35] were developed for model assessment. The 10 largest variable importance measures from the selected base models of the augmented experiment were also compared [15, 36]. Model assessment was performed using the boot [37, 38], Metrics [39], and ResourceSelection [40] packages in the R statistical software. Variable importance measures were gathered using the h2o package [13].

## 3. Results

### 3.1. Model Performance and Variable Importance

The performance metrics of each model from the ICD-9 experiment (where only information pertaining to ICD-9 codes in the processed data set was considered as input variables) are displayed in Table 3. For the ICD-9 experiment, TSM demonstrates an improvement over BLISS, HARM, and TMPM for each performance metric. TSM also demonstrates an improvement over its base models for nearly every performance metric (the selected random forest model has better HL). The performance metrics of each model from the augmented experiment are displayed in Table 4. Every model greatly improved in performance when augmented to account for patient demographics (age and gender) as well as general trauma assessments (comorbidities, Glasgow Coma Scale response scores, injury mechanism, injury type, and intent of trauma). But, TSM outperforms every other model for each performance metric. No single base model in TSM’s ensemble consistently outperforms all other base models for every performance metric in each experiment. BLISS outperforms TSM’s base models for most performance metrics in each experiment.

**Table 3:**
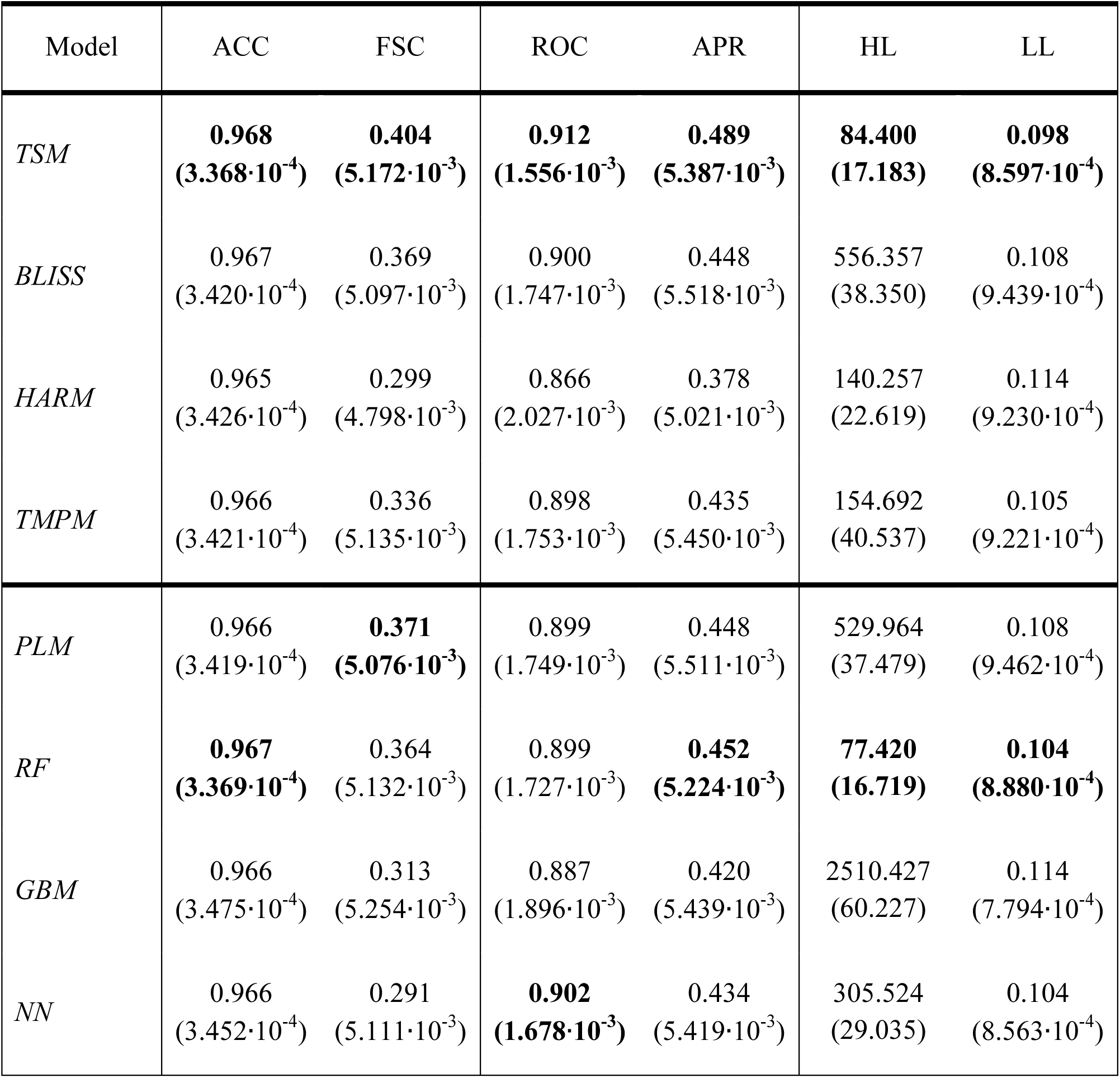
Model comparison for the ICD-9 experiment. Standard error of the metric is denoted in the parenthesis. TSM consistently has superior performance under each performance metric except for the HL statistic, which was attained by the random forest base model. PLM, RF, GBM, and NN denote logistic regression with the elastic net penalty, random forest, gradient boosted machine, and neural network, respectively.

**Table 4:**
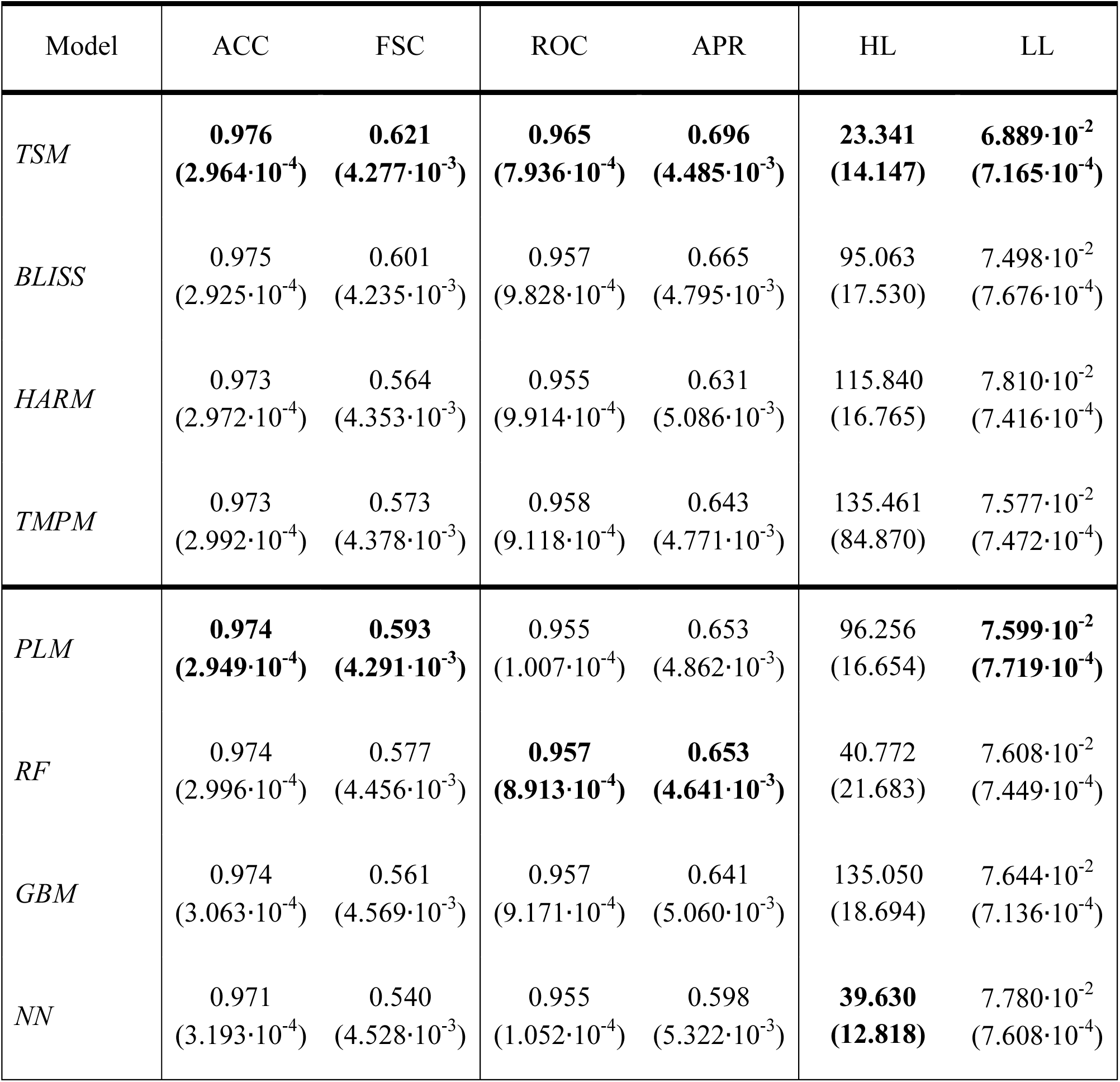
Model performance comparison for the augmented experiment. Standard error of the metric is denoted in the parenthesis. TSM consistently has superior performance under each metric. PLM, RF, GBM, and NN denote logistic regression with the elastic net penalty, random forest, gradient boosted machine, and neural network, respectively.

The 10 largest variable importance measures from the selected base models of the augmented experiment are displayed in Figure 1. Each model ranks the significance of their input variables differently. But, a Glasgow Coma Scale eye response score of 1 and a patient’s age were amongst the 10 largest variable important measures for all selected base models. In general, each selected base models heavily relies on information pertaining to head trauma at the time of admission when predicting patient outcomes. The models differed in that the selected logistic regression model developed with the elastic net penalty placed a large variable importance measure on neck sprains; the random forest placed a large variable importance measure on congestive heart failure; the gradient boosted machine placed a large variable importance measure on lung injury; and the neural network placed a large variable importance measure on being physically struck by a person or object.

**Figure 1:**
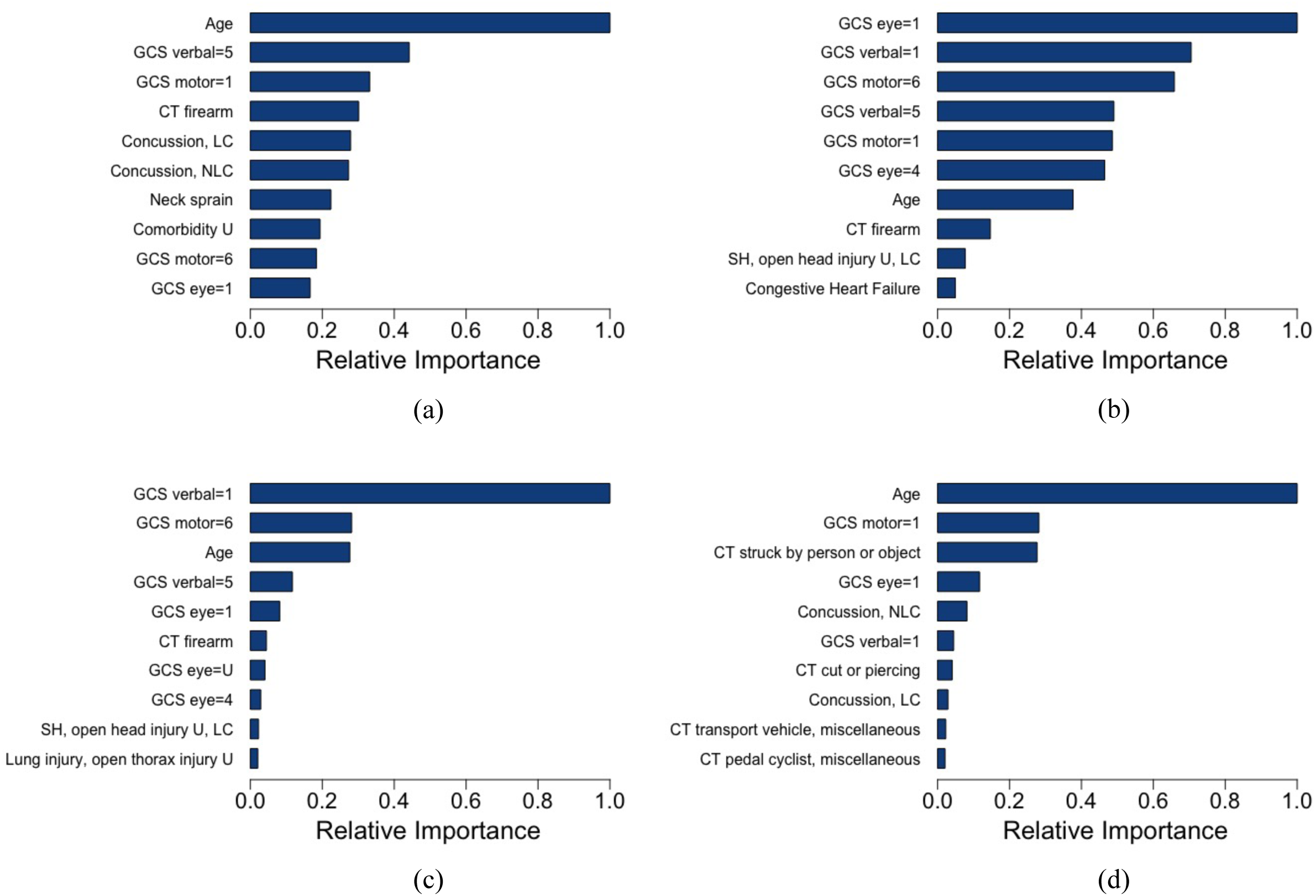
10 largest variable importance measures of the selected base models from TSM’s ensemble for the augmented experiment. The selected logistic regression model developed with the elastic net penalty is denoted by (a), random forest by (b), gradient boosted machine by (c), and neural network by (d). Further, GCS denotes Glasgow Coma Scale, CT denotes cause of trauma, LC denotes loss of consciousness, NLC denotes no loss of consciousness, U denotes unknown, and SH denotes subdural hemorrhage.

The calibration curves of TSM, BLISS, HARM, and TMPM models from the ICD-9 experiment are displayed in Figure 2; the calibration curves of TSM, BLISS, HARM, and TMPM models from the augmented experiment are displayed in Figure 3. TSM and BLISS consistently provide well-calibrated prediction outputs, whereas TMPM and HARM do not provide well-calibrated prediction outputs for the ICD-9 experiment. The precision-recall curves of TSM, BLISS, HARM, and TMPM models from the ICD-9 experiment are displayed in Figure 4; the precision-recall curves of TSM, BLISS, HARM, and TMPM models from the augmented experiment are displayed in Figure 5. TSM generally displays higher precision and recall than BLISS, HARM, and TMPM for all thresholds in both experiments.

**Figure 2:**
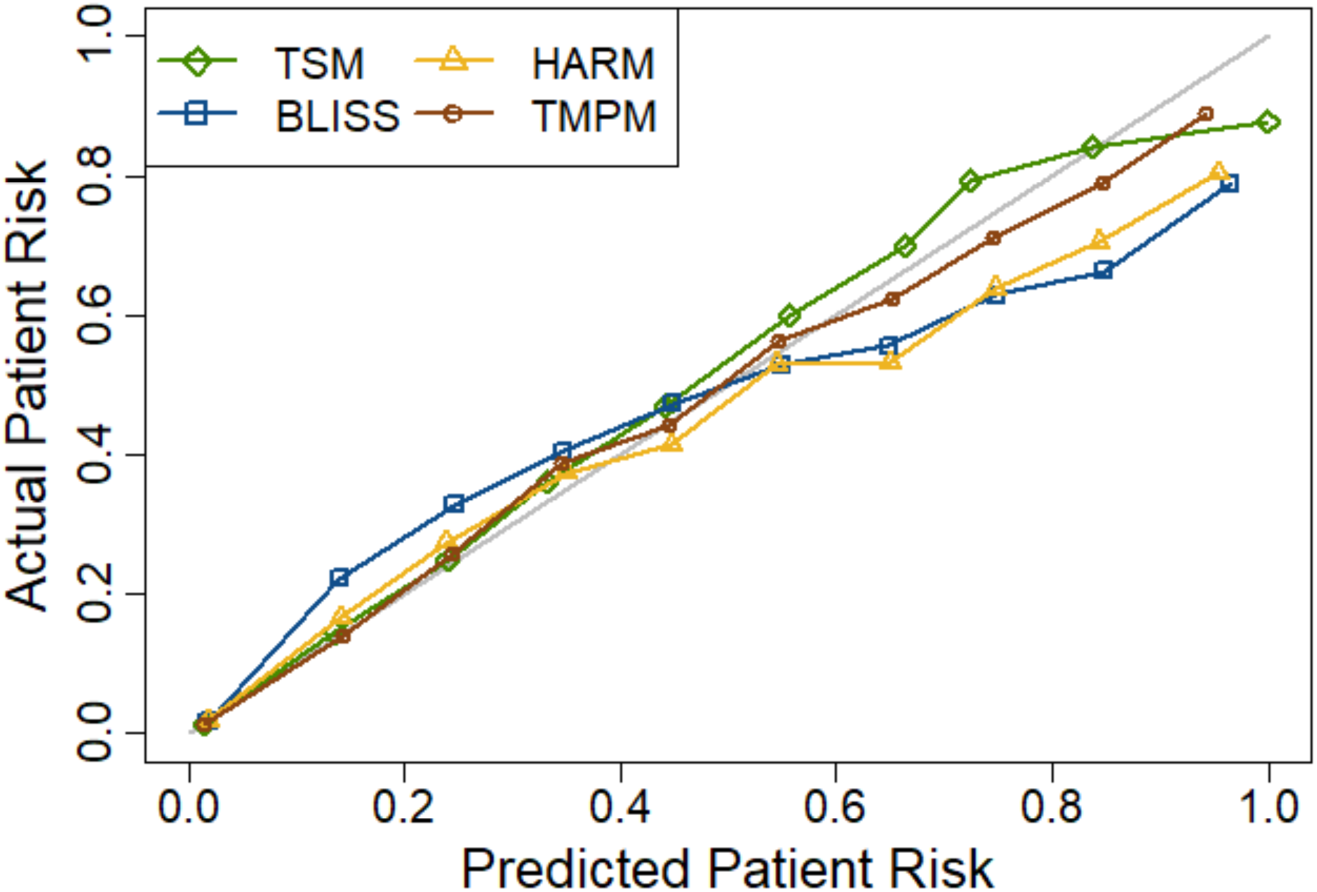
Calibration curves for TSM, HARM, BLISS, and TMPM models from the ICD-9 experiment. The grey line represents perfect probabilistic calibration. TSM and TMPM provide well-calibrated prediction outputs.

**Figure 3:**
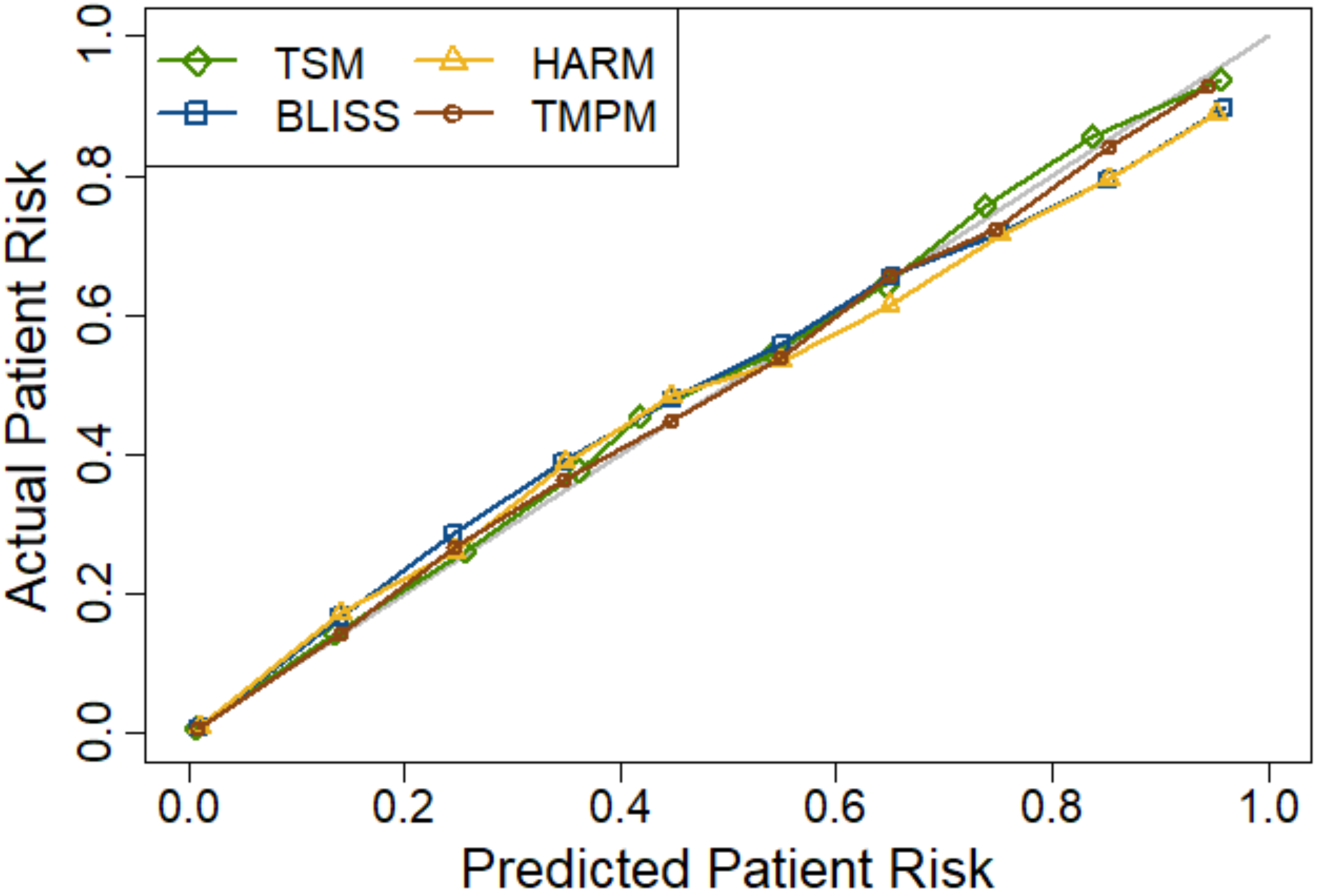
Calibration curves for TSM, HARM, BLISS, and TMPM models from the augmented experiment. The grey line represents perfect probabilistic calibration. Every model provides well-calibrated prediction outputs.

**Figure 4:**
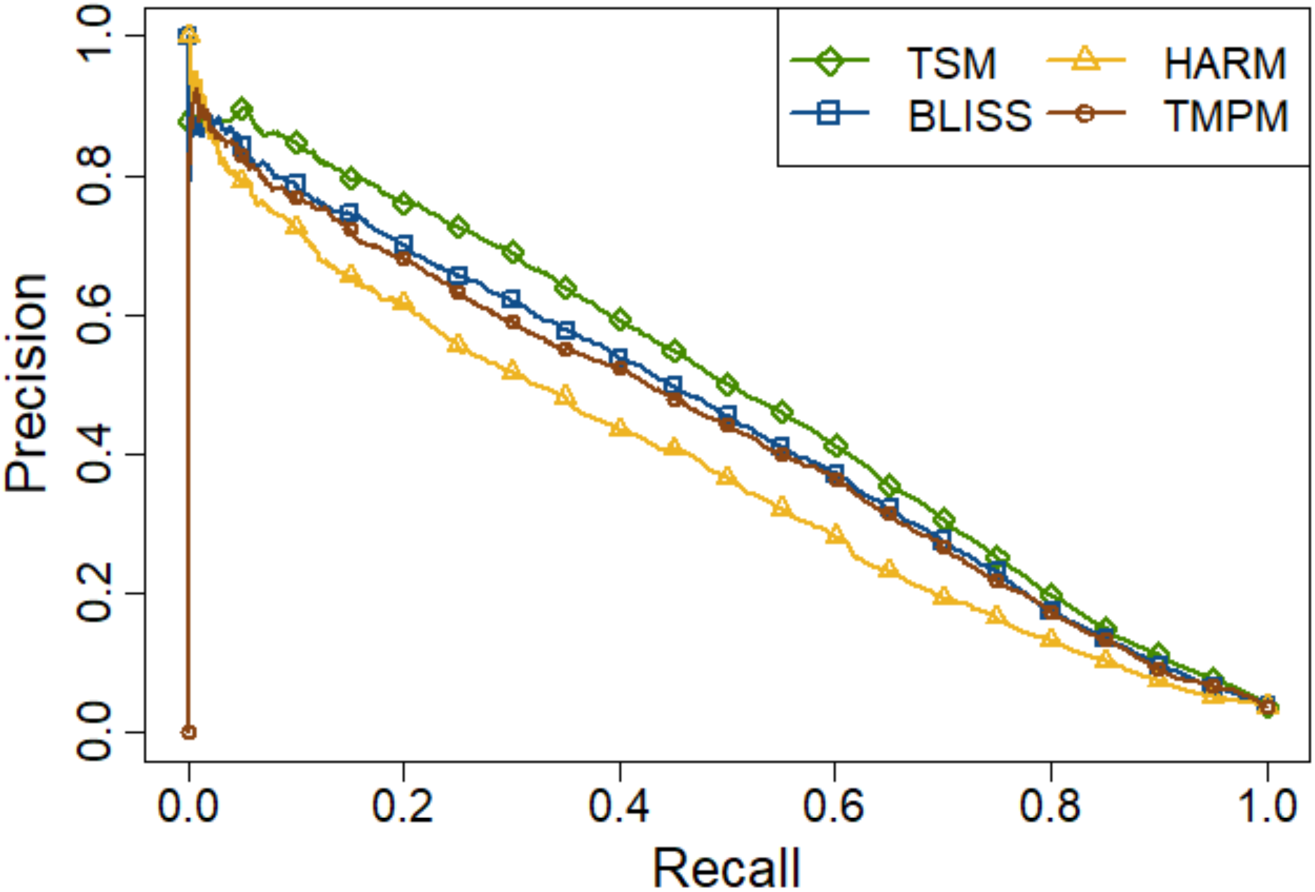
Precision-recall curves for TSM, BLISS, HARM, and TMPM models from the ICD-9 experiment. TSM generally had higher precision and recall for all thresholds.

**Figure 5:**
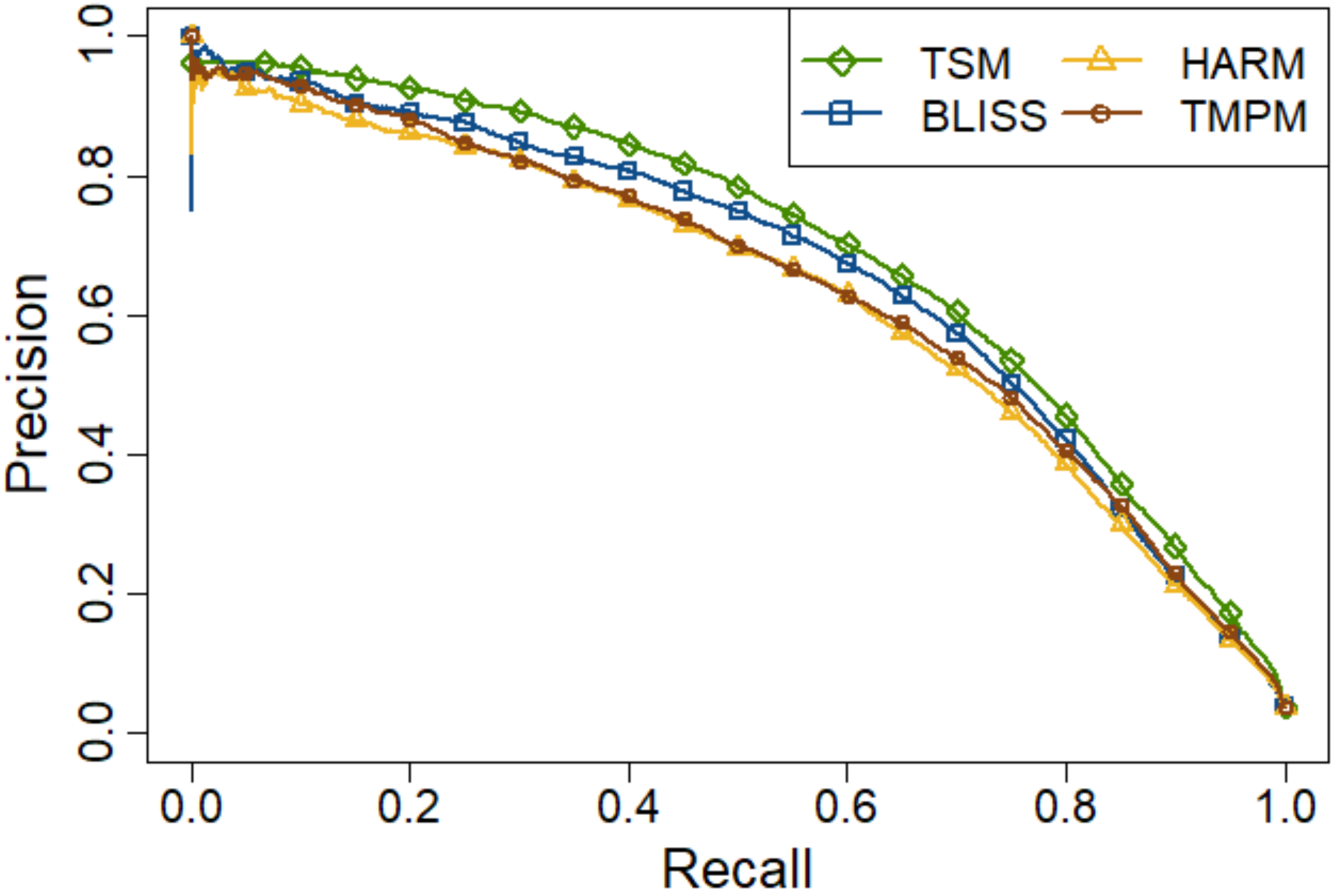
Precision-recall curves for TSM, BLISS, HARM, and TMPM models from the augmented experiment. TSM generally had higher precision and recall for all thresholds.

Figure 6 shows the performance of our hyper-parameter search scheme for the ICD-9 experiment, and Figure 7 shows the performance of our hyper-parameter search scheme for the augmented experiment. Figure 6 demonstrates that our hyper-parameter search scheme was particularly successful when developing random forest models, as hyper-parameter space shifts correspond to decreasing LL on the validation set. Figure 7 shows that no hyper-parameter space shifting occurred during this experiment. Table 5 displays the hyper-parameters of the selected machine learning base models from each experiment (the models with lowest LL on the validation set). The maximum tree depth hyper-parameter of the meta-learner selected for TSM was 1 in both experiments.

**Figure 6:**
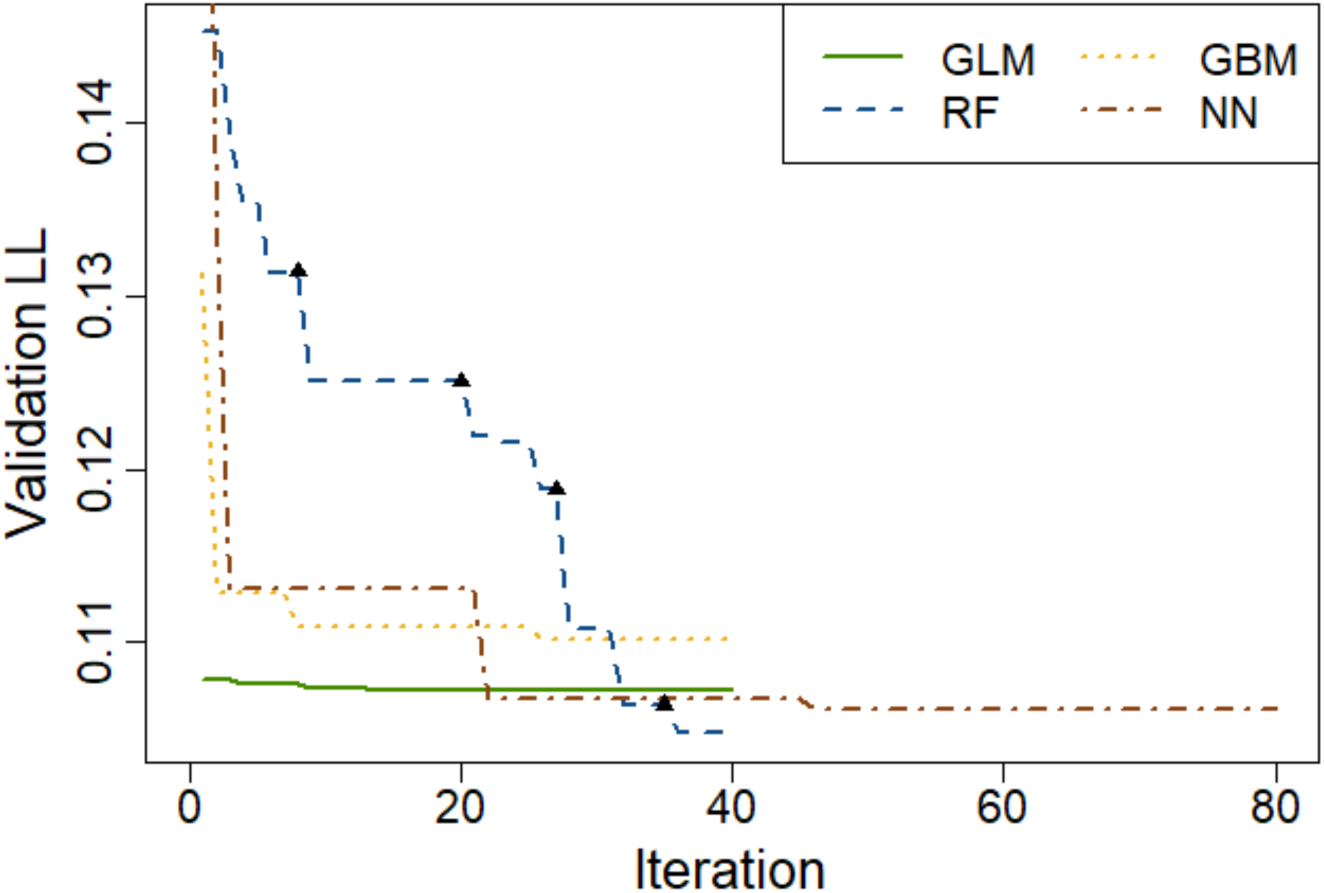
Log-loss on the validation set (validation LL) for the model that had the lowest validation LL after each iteration of our hyper-parameter search procedure from the ICD-9 experiment. The hyper-parameter space shifted four times for the random forest algorithm, and each shift is associated with a decrease in validation LL. Shifts are denoted by a black triangle (▲).

**Figure 7:**
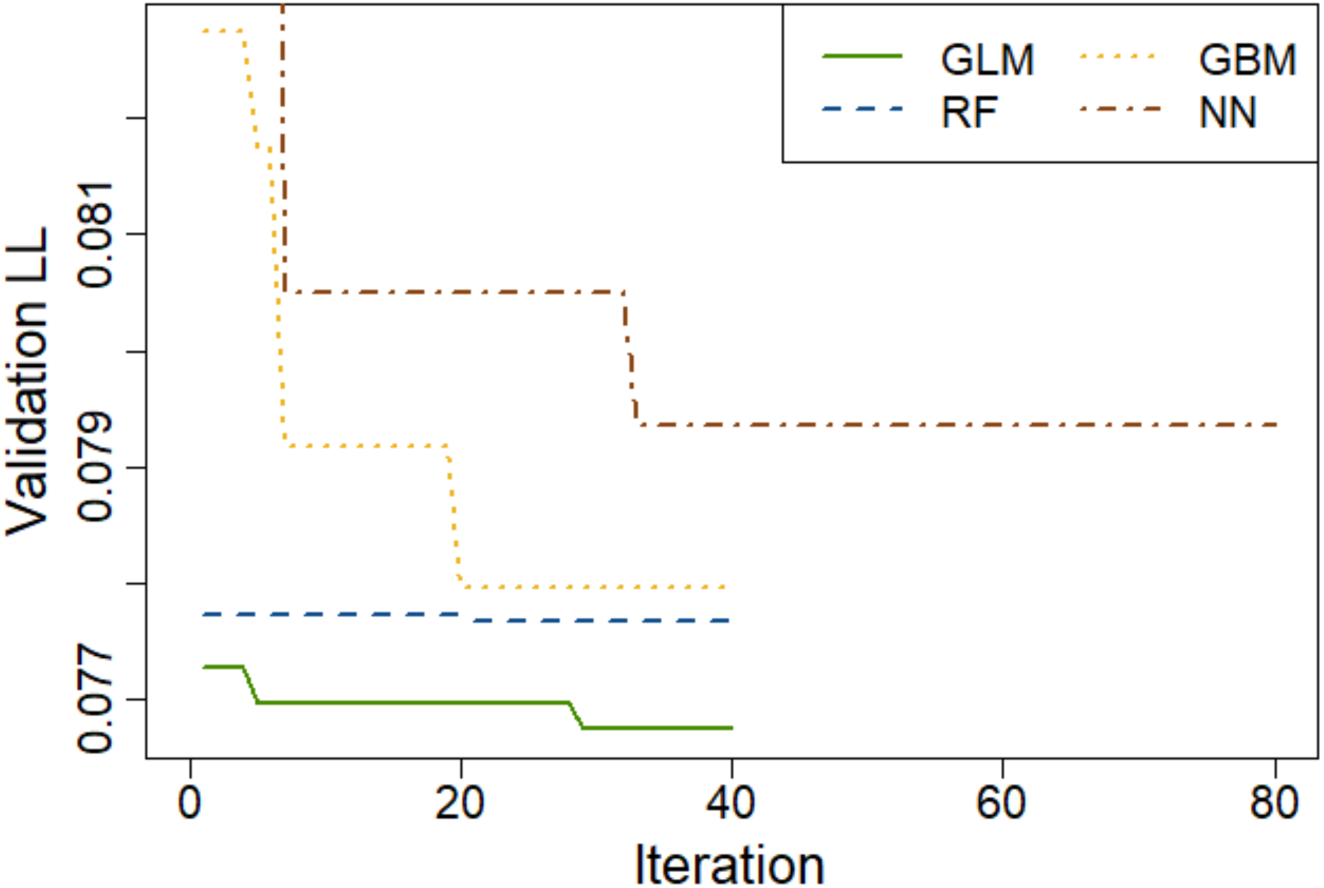
Log-loss on the validation set (validation LL) for the model that had the lowest validation LL after each iteration of our hyper-parameter search procedure from the augmented experiment. The hyper-parameter space did not shift for any hyper-parameter search.

**Table 5:**
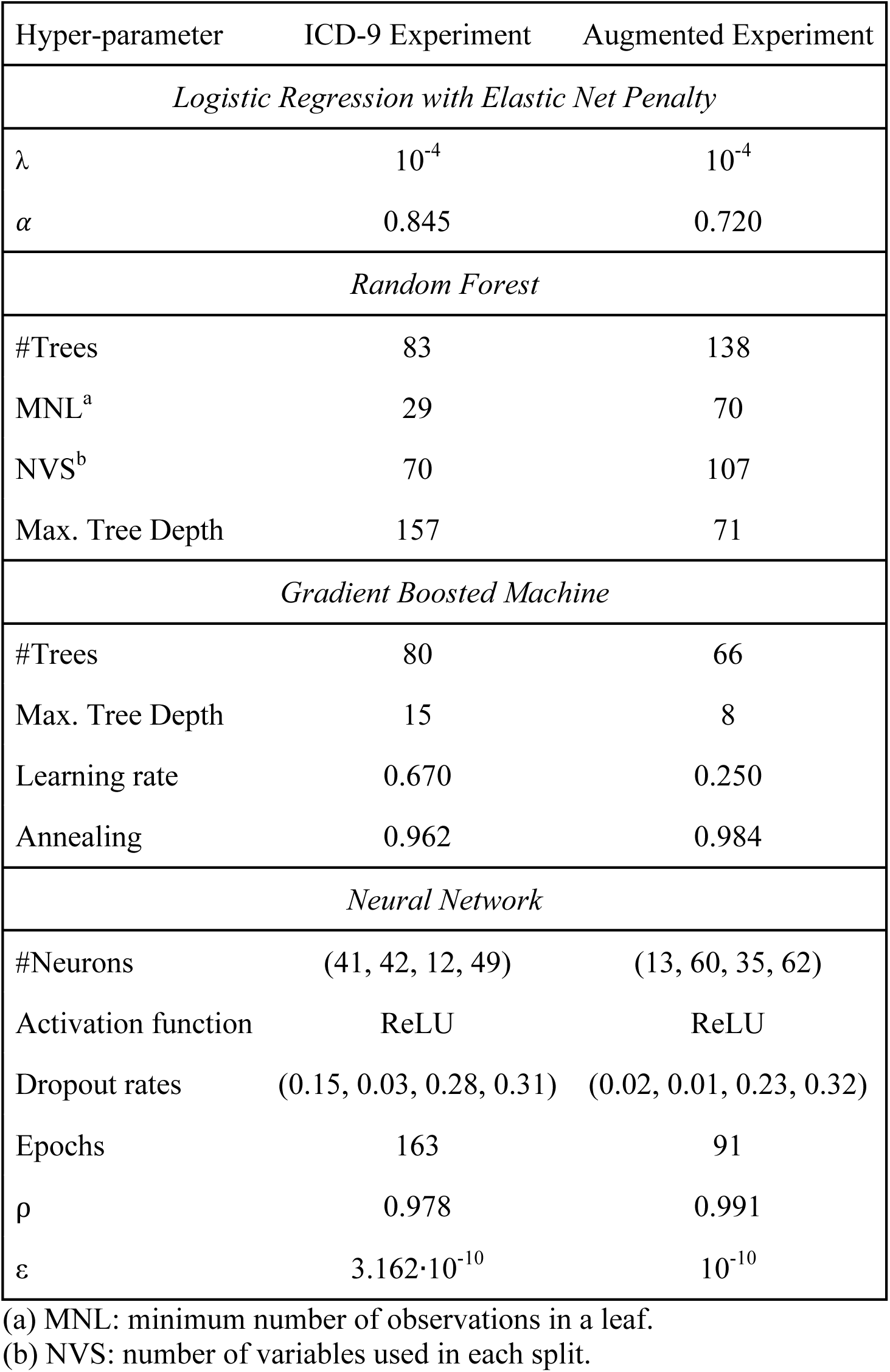
Hyper-parameters of the selected base models from each experiment.

### 3.2. Discussion

Trauma is the leading cause of death for people younger than 44, and the fourth leading cause of death for all age groups in the United States [41]. As healthcare spending has grown to 17.8% of the Gross Domestic Product, it is increasingly important to take the cost of care into consideration when improving the quality of trauma care [42]. But in order to achieve the goals of improving the quality of trauma care while controlling the cost of care, we must utilize the best possible risk prediction models in trauma system evaluations. If risk prediction can be improved, so too can the quality of trauma care, as better risk prediction models allow for a better evaluation of novel treatments, interventions, and policies. This study demonstrates that TSM, a machine learning model, outperforms established risk prediction on every performance metric considered in this study.

There is controversy regarding the utility of machine learning in healthcare [43]. This is in part motivated by several studies that compared generalized linear models to individual machine learning models for risk prediction, often with contradictory results [44-47]. This phenomenon is in part due to the fact that no single algorithm is inherently better than all others – depending on the performance metric and the complexity of the data, the best predictive model may be developed from any algorithm [48, 49]. This claim is evidenced by the results of this study, as TSM’s base models outperform established risk prediction models on some performance metrics, while established risk prediction models outperform TSM’s base models on other performance metrics.

What separates an ensemble machine learning approach, such as stacked generalization, from a methodology where a single model is selected and assessed is that, if performed appropriately, stacked generalization will utilize its base models’ strengths while compensating for their weaknesses. As a result, it is likely that a well-designed ensemble machine learning model developed from stacked generalization will obtain better predictive performance than any base model in its ensemble [17, 18, 50-52]. This is also indicated by the results in this study, as TSM outperforms its base models on nearly every performance metric.

The challenge with developing a well-designed ensemble machine learning model is that the ensemble must consist of base models that have strong predictive performance (ensemble strength) as well as base models that provide different prediction outputs for the same conditions (ensemble diversity). Our hyper-parameter search scheme attempts to address both of these, as a random search can provide both ensemble strength and ensemble diversity with regards to a hyper-parameter space, and hyper-parameter space shifting attempts to improve hyper-parameter space configuration if the optimal hyper-parameters lie beyond the initial hyper-parameter space configured.

While our hyper-parameter search scheme worked for this study, our search scheme is not guaranteed to work for all settings, as it is dependent on the sensitivity of the initial hyper-parameter space configured as well as the state of the random number generator. Issues pertaining to sensitivity may be addressed by taking careful measures to configure an appropriate initial hyper-parameter space. Issues pertaining to random number generation may be addressed by developing a large number of models, examining a large number of models with regards to a small portion of each hyper-parameter interval, and specifying a small distance to shift a hyper-parameter interval.

A potential concern with the results of this study is that our hyper-parameter search procedure may have been insufficient due to BLISS outperforming the non-linear base models on most performance metrics. But, this is a consequence of re-calibrating these non-linear models. In particular, due to the tradeoff between discrimination and probabilistic calibration [53], the ROC of the random forest, gradient boosted machine, and neural network base models selected for model assessment diminished. Although this improved the probabilistic calibration of these models, some models, such as the selected gradient boosted machine in the ICD-9 experiment, were so poorly calibrated that their overall performance appeared poor when re-calibrated. While this study highlights the strengths of developing an ensemble machine learning model, most medical studies do not require anything more sophisticated than a generalized linear model. This is due to their low computational cost, their simple functional form (which captures the underlying relationships in most medical data sets), and the interpretability as well as consistency of their weights. Arguably, generalized linear models could still be considered most appropriate for mortality risk prediction with trauma patients, as the established risk prediction models have strong predictive performance, and it is currently unknown whether or not TSM will display a clinically significant improvement over these established risk prediction models in other settings.

However, as modeling problems with healthcare data become increasingly complex, non-linear machine learning algorithms should be considered, as they can automatically find non-linear relationships in data [16]. Generalized linear models, on the other hand, would require extensive feature engineering for such data, and depending on the setting such measures will not result in the development of a model that performs as well as a model developed from machine learning algorithms. This claim is partly validated by our results, as HARM, which is developed using extensive feature engineering based on clinical intuition and expert consensus, generally had worse performance metrics than TSM’s base models. Further, the variable importance measures of the non-linear models from Figure 1 reflect reality, as firearm injuries (ranked highly by the random forest) as well as vehicular accidents (ranked highly by the neural network) are considered significant variables in predicting patient outcome [11]. This indicates that the variable importance measures of non-linear machine learning models can also have value in studies necessitating interpretability.

To address a major concern with the use of machine learning algorithms in healthcare, these results do not imply that the prediction outputs of machine learning models should replace expert opinion. But, the use of machine learning models with expert opinion can greatly improve patient care in a variety of settings, as indicated in [54].

## 4. Conclusions

The Trauma Severity Model improves over established risk prediction models for every performance metric considered in this study, which gives it prognostic value in trauma system evaluations. The hyper-parameter search scheme proposed for this study performed well and developed strong performing machine learning models. The performance of an ensemble machine learning model on a well-studied problem in epidemiology indicates that ensemble machine learning approaches may be fruitful for other complex problems in healthcare.

## Conflict of Interest

The authors declare that they have no conflicts of interest in regards to the content in this article.

## Acknowledgements

The authors would like to thank Dr. Laurent Glance and Dr. Turner Osler for their invaluable assistance on this project. The authors would also like to thank Ph.D. student Hugo Milan for conversations concerning this manuscript. These results were presented at the Eastern Association for the Surgery of Trauma Annual Scientific Assembly in 2017.

## Summary

Previous studies on trauma mortality prediction from ICD-9 codes (800-959.9) have focused on the use of generalized linear models for risk prediction. Although this has resulted in beneficial mortality prediction models, there are a variety of other algorithms that may lead to the development of a model with even better predictive performance. In this study, we have developed several predictive models from different machine learning algorithms, and we have compared their predictive performance to the performance of established, widely used trauma risk prediction models. Our results indicate that these individual machine learning models have comparable performance to the established trauma risk prediction models. However, combining these machine learning models into an ensemble (using stacked generalization) leads to the development of a model with better predictive performance than the established trauma risk prediction models for each performance metric considered. Previously, stacked generalization has seldom been considered in this setting. This study indicates that intensive data-driven approaches can improve our ability to predict mortality risk of trauma patients, and thereby delineate patients in need of aggressive care.

